# Monkey dorsolateral prefrontal cortex represents abstract visual sequences during a no-report task

**DOI:** 10.1101/2022.09.19.508576

**Authors:** Nadira Yusif Rodriguez, Theresa H. McKim, Debaleena Basu, Aarit Ahuja, Theresa M. Desrochers

## Abstract

Monitoring sequential information is an essential component of our daily lives. Many of these sequences are abstract, in that they do not depend on the individual stimuli, but do depend on an ordered set of rules (e.g., chop then stir when cooking). Despite the ubiquity and utility of abstract sequential monitoring, little is known about its neural mechanisms. Human rostrolateral prefrontal cortex (RLPFC) exhibits specific increases in neural activity (i.e., “ramping”) during abstract sequences. Monkey dorsolateral prefrontal cortex (DLPFC) has been shown to represent sequential information in motor (not abstract) sequence tasks, and contains a sub-region, area 46, with homologous functional connectivity to human RLPFC. To test the prediction that area 46 may represent abstract sequence information, and do so with parallel dynamics to those found in humans, we conducted functional magnetic resonance imaging (fMRI) in monkeys. When monkeys performed no-report abstract sequence viewing, we found that left and right area 46 responded to abstract sequential changes. Interestingly, responses to rule and number changes overlapped in right area 46 and left area 46 exhibited responses to abstract sequence rules with changes in ramping activation, similar to that observed in humans. Together, these results indicate that monkey DLPFC monitors abstract visual sequential information, potentially with a preference for different dynamics in the two hemispheres. More generally, these results show that abstract sequences are represented in functionally homologous regions across monkeys and humans.

**Significance Statement:** Daily, we complete sequences that are “abstract” because they depend on an ordered set of rules (e.g., chop then stir when cooking) rather than the identity of individual items. Little is known about how the brain tracks, or monitors, this abstract sequential information. Based on previous human work showing abstract sequence related dynamics in an analogous area, we tested if monkey dorsolateral prefrontal cortex (DLPFC), specifically area 46, represents abstract sequential information using awake monkey fMRI. We found that area 46 responded to abstract sequence changes, with a preference for more general responses on the right and dynamics similar to humans on the left. These results suggest that abstract sequences are represented in functionally homologous regions across monkeys and humans.

## Introduction

Sequential tasks that require monitoring are prevalent in daily life. For example, taking a bus requires tracking familiar sequences of buildings (e.g., three houses then a library), enabling the detection of deviations from this sequence (e.g., if there is a detour). Similarly, many cognitive processes occur serially, and often demand that we maintain an internal representation of the previous steps to complete the next. Even in tasks that are not explicitly sequential, a system for tracking transitions between steps, such as when completing a mathematical operation, may be essential.

*Sequence monitoring* is this active process of tracking the order of subsequent “states” or steps. Monitoring is distinct from other well-studied sequence processes, such as explicit memorization, or potentially more automatic behaviors, such as a series of motor outputs (e.g., playing the piano) or statistical sequence learning (Desrochers et al., 2019). Such sequential processes likely contain monitoring operations within them but are also comprised of other cognitive computations. *Abstract sequences* are sequences that are not dependent on the individual stimuli but can instead be described by the rule they follow (e.g., three same, one different or AAAB) (Desrochers et al., 2022). Therefore, *abstract sequence monitoring* entails sequences of sensory stimuli that possess abstract structure and active monitoring of this structure. While it may be assumed that abstract sequence monitoring underlies many aforementioned sequence types (including motor sequences), it is rarely studied in isolation and the neural underpinnings of abstract sequence monitoring remain largely unknown.

Multiple modes of evidence in humans indicate that activity in rostrolateral prefrontal cortex (RLPFC) is crucial to sequence monitoring (Desrochers et al., 2015, 2019; McKim and Desrochers, 2022). Functional magnetic resonance imaging (fMRI) revealed systematic increasing activity (“ramping”) from the beginning to the end of each sequence in human RLPFC. Across studies, this activity occurs either bilaterally, or in the left RLPFC. Further, noninvasive transcranial magnetic stimulation (TMS) showed that the left RLPFC was necessary for sequential tasks in humans. Other studies have also demonstrated the involvement of RLPFC as part of a frontoparietal network active during complex sequential tasks (Farooqui et al., 2012; Wang et al., 2019; Wen et al., 2020). While consistent with the findings discussed above, these studies frequently involve other cognitive phenomena, like decision-making, leaving open their specific role of sequence monitoring.

Studies in nonhuman primates also suggest a role of lateral prefrontal cortex in abstract visual sequence monitoring. Motor sequence studies show that neurons in the dorsolateral prefrontal cortex (DLPFC) are selective for serial position (Barone and Joseph, 1989; Averbeck et al., 2006; Shima et al., 2007; Berdyyeva and Olson, 2010) and sequence boundaries (Fujii and Graybiel, 2003), and include neural dynamics that could underlie the ramping observed in human BOLD activation(Desrochers et al., 2015, 2019; McKim and Desrochers, 2022). Neurons in the DLPFC also show ordinal selectivity during visual object sequences (Ninokura et al., 2004; Warden and Miller, 2010; Naya et al., 2017). A rich literature also supports the involvement of DLPFC in representing non-sequential abstract rules (Hoshi et al., 1998; White and Wise, 1999; Wallis et al., 2001; Eiselt and Nieder, 2013). Responses in the DLPFC can also selectively represent sequential regularities (Vergnieux and Vogels, 2020). Together, these physiological studies suggest that the monkey DLPFC is well-positioned to monitor abstract visual sequences. We hypothesized that a specific sub-region of monkey DLPFC (area 46) monitors visual abstract sequential information. In humans, abstract sequence monitoring has been localized to the RLPFC, which is distinct from the rostromedial prefrontal cortex (Koechlin et al., 2000; Burgess et al., 2003; Gilbert et al., 2010; Moayedi et al., 2015; Henssen et al., 2016; Du et al., 2020). While rostromedial prefrontal cortex has similar connectivity in monkeys and humans, anatomical evidence suggests that monkey area 46 contains the most similar connectivity patterns to human RLPFC (Sallet et al., 2013; Neubert et al., 2014), and overlapping high-level visual representations (Xu et al., 2022). Therefore, we predicted that abstract visual sequence monitoring would be supported by monkey area 46, and that similar ramping dynamics, as observed in humans, would localize to this same area.

To directly test these predictions, we conducted event-related fMRI in awake nonhuman primates, and used deviations from established abstract visual sequences during a no-report task to index abstract sequence monitoring. We found that nonhuman primate DLPFC distinctly represents abstract visual sequence information, independent from other task constraints. Additionally, we find that these abstract sequences elicit ramping dynamics similar to those observed in humans during abstract sequence performance. Intriguingly, deviant responses with differing primary dynamics were observed in the two hemispheres: an onset-based signal on the right, and ramping on the left. These findings indicate that a specific sub-region of DLPFC preferentially supports abstract sequence monitoring in monkeys and may be functionally homologous to humans. Further, these results establish an important connection between human and monkey complex cognition and the neural substrates that mediate it, providing a foundation for understanding more complex behaviors across species in the future.

## Materials and Methods

### Subjects

We tested three adult male rhesus macaques (ages spanning 6-12 years during data collection, 9-14 kg). All procedures followed the NIH Guide for Care and Use of Laboratory Animals and were approved by Institutional Animal Care and Use Committee (IACUC) at Brown University.

### No-Report Abstract Visual Sequence Task

All visual stimuli used in this study were displayed using an OpenGL-based software system developed by Dr. David Sheinberg at Brown University. The experimental task was controlled by a QNX real-time operating system using a state machine. Eye position was monitored using video eye tracking (Eyelink 1000, SR Research). Stimuli were displayed at the scanner on a 24-inch BOLDscreen flat-panel display (Cambridge Systems). The general design of the visual sequence paradigm was based on a similar auditory sequence task (Wang et al., 2015).

#### Stimuli

Each image presentation consisted of fractal stimulus (approximately 8° visual angle) with varying colors and features. Fractals were generated using MATLAB for each scanning session using custom scripts based on stimuli from (Kim and Hikosaka, 2013) following the instructions outlined in (Miyashita et al., 1991). For each scan session, new, luminance matched, fractal sets were generated. All stimuli were presented on a gray background, with a fixation spot that was always present on the screen superimposed on the images. To provide behavioral feedback, the fixation spot was yellow when the monkey was successfully maintaining fixation and red if the monkey was not fixating. Stimuli were displayed for 0.1 to 0.3 s each, depending on the sequence type and timing template, detailed as follows.

#### Sequence Types

There are five sequence types in this task (**Figure 1**): habituation sequences and four deviant sequence types. Across these sequence types, there were a total of nine different timing templates used. These templates were included to counterbalance for stimulus and sequence duration across the sequence types and to provide a greater variety of sequential timings during habituation. The inter-sequence interval was jittered to decorrelate across timing templates (mean 2 s, 0.25-8 s).

**Figure 1.**
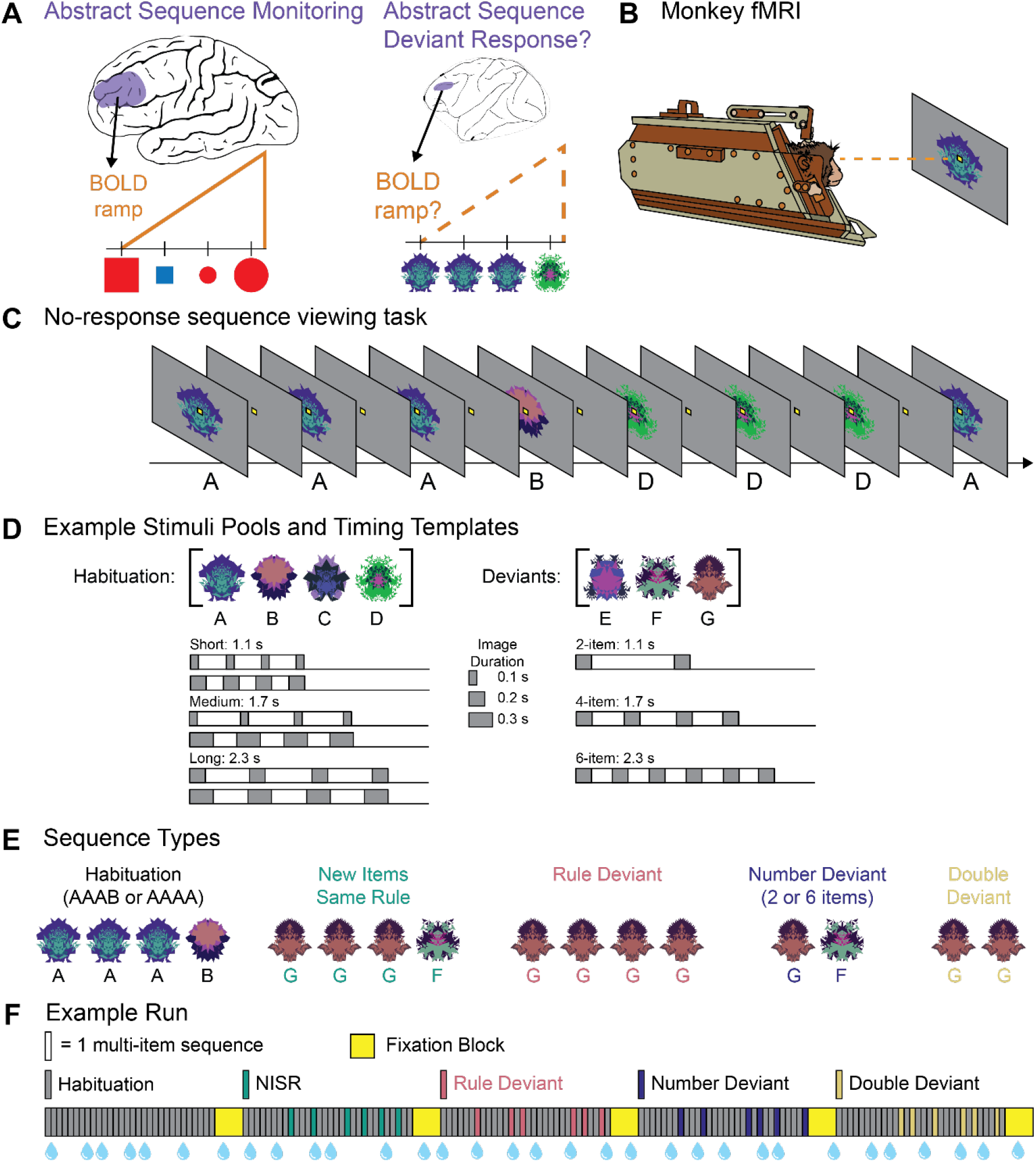
No-report abstract sequence viewing task. **A**. Schematic representation of human rostrolateral prefrontal cortex (RLPFC; left) and monkey dorsolateral prefrontal cortex (DLPFC; right) depicting the main questions that were the focus of this study: Does monkey DLPFC monitor abstract sequences, as shown by responses to deviant sequences? and does monkey DLPFC exhibit ramping activation, as found in human RLPFC during sequence monitoring? **B**. Monkeys only fixate throughout runs. Scanning is performed in the “sphynx” position. **C**. Example partial habituation block for sequence rule *three same, one different* (AAAB). **D**. Example stimulus pools (top) show a set of images that would be used in a single scanning session. New images are used each session. Six possible timing templates for habituation sequences (bottom, left) and deviant sequences (bottom, right) illustrated with gray rectangles indicating single images. Total sequence durations are listed for each template type. **E**. Examples of the five sequence types if the sequence rule in use is *three same, one different*. **F**. Example run, with each bar indicating one multi-image sequence: four images in habituation, new items same rule (NISR), and rule deviants; two or six images in number and double deviants. The first block contains only habituation sequences and subsequent blocks contain only one of the four deviant types. Sequence blocks alternate with fixation blocks. Blue water droplets schematize reward delivery, which is decoupled from sequence viewing and delivered on a graduated schedule based on the duration the monkey has maintained fixation.

##### Habituation Sequences

Habituation sequences were composed of images drawn from a pool of four possible fractals. We will refer to the habituation image pool as [A, B, C, D]. Sequences were composed from these images in one of two possible rules: three the same, one different (e.g., AAAB, DDDC) and four the same (e.g., AAAA, CCCC). All sequences contained four images and followed one of three possible general timings based on the total duration of the sequence: short (1.1 s), medium (1.7 s), and long (2.3 s). Each total sequence duration, in turn, had two possible timing templates within it, one with longer stimulus durations and one with shorter stimulus durations: short 0.1 s and 0.2 s, medium 0.1 s and 0.3 s, long 0.2 s and 0.3 s. Inter-stimulus intervals were arranged to evenly space the four stimulus presentations within the total sequence duration.

##### Deviant Sequences

Deviant sequences were composed of images drawn from a different pool of three possible fractals. We will refer to the deviant image pool as [E, F, G]. All deviant images were displayed for 0.2 s, regardless of deviant type. Across deviant types, the total sequence durations were matched to the short, medium, and long habituation timing templates. There were four deviant types, detailed as follows:

##### New Items, Same Rule (NISR)

These deviants use images that come from the deviant pool of images, but do not differ from the habituation rule. For example, if the habituation rule was three the same, one different then NISR sequences would follow the same rule with new images (e.g., GGGF and FFFE) Alternatively, if the habituation rule was four same, an example NISR would be EEEE. All sequences were four items and had a total duration of 1.7 s.

##### Rule deviants

These deviants do not follow the same rule as habituation, but instead follow the alternate rule. If the habituation rule was three the same, one different, example deviants would follow the four the same rule, e.g., EEEE and GGGG. All deviants contained four images and had a total sequence duration of 1.7 s, the same as medium habituation sequences.

##### Number Deviants

These deviants follow the same rule as habituation but contain a different number of images (either two or six). If the habituation rule was three the same, one different, example deviants would be EG and FFFFFE. Two-item deviants had a total sequence duration of 1.1 s, the same as short habituation sequences, and six-item deviants had a total sequence duration of 2.3 s, the same as long habituation sequences.

##### Double Deviants

These deviants combine Rule and Number deviant types. If the habituation rule was three the same, one different, example deviants would be EE and GGGGGG. The timing was the same as number deviants.

#### Block Structure

Each block contained 30 sequences and lasted approximately 112 s on average. Habituation blocks contained equal numbers of the six possible timing templates (two of each: short, medium, and long). Habituation sequences were presented in pseudo-random order such that a sequence could not begin with the same fractal as the final fractal of the previous sequence. Deviant blocks were composed of 24 habituation sequences and six deviant sequences. All deviant sequences within a block were of the same sequence type. The six deviant sequences were pseudo-randomly interspersed throughout the block such that deviant sequences did not occur in the first 6 sequences of the block (to avoid block initiation confounds), and deviant sequences were not presented consecutively to each other. If deviant sequences contained a variable number of items (i.e., number deviants and double deviants), then an equal number of two- and six-item sequences were included within a single block. The 24 habituation sequences within deviant blocks were presented in the same manner as in habituation blocks (i.e., evenly distributed timing templates and avoiding between-sequence fractal image repeats).

#### Run Structure

Each run was composed of five blocks, interleaved with 14 s fixation blocks (**Figure 1**). The first block of each run contained only habituation sequences. The four subsequent blocks were one of each of the four possible deviant types, with their order counterbalanced across runs. The same habituation rule was used for the entirety of a single run. Runs lasted approximately 10.5 min. The sequence rule (*three same, one different* or *four same*) used for each run was counterbalanced across each scanning session so as to have an equal number of runs for each rule. Monkeys typically completed 4-8 runs of this task (among other tasks not reported on here) in a single scanning session (one day).

Runs were initiated according to the monkey’s fixation behavior to ensure that the monkey was not moving and engaged in the task before acquiring functional images. During this pre-scan period, a fixation spot was presented. Once the monkey successfully acquired this fixation spot and received approximately four liquid rewards (12 – 16 s), functional image acquisition and the first habituation block were initiated.

#### Reward

The timing of liquid rewards was not contingent upon sequential events, only on the monkey maintaining fixation. Rewards were delivered on a graduated schedule such that the longer the monkey maintained fixation, the more frequent rewards were administered (Leite et al., 2002). The first reward was given after 4 s of continuous fixation. After two consecutive rewards of the same fixation duration, the fixation duration required to obtain reward was decreased by 0.5 s. The minimum duration between rewards that the monkey could obtain was 0.5 s. Fixation had to be maintained within a small window (typically 3° of visual angle) around the fixation spot to not break fixation. The only exception was a brief time window (0.32 s) provided for blinks. If the monkey’s eyes left the fixation window and returned within that time window, it would not trigger a fixation break. If fixation was broken, the reward schedule would restart at the maximum 4 s duration required to obtain reward.

### FMRI Data Acquisition

Monkeys were trained to sit in the “sphynx” position in a custom MR-safe primate chair (Applied Prototype, Franklin, MA or custom-made by Brown University). The monkey’s head was restrained from moving via a plastic “post” (PEEK, Applied Prototype, Franklin, MA) affixed to the monkeys’ head and the primate chair. Monkeys were habituated to contrast agent injection procedures, recorded MRI sounds, wearing earplugs (Mack’s Soft Moldable Silicone Putty Ear Plugs, Kid’s size), and transportation to the scanner prior to MRI scanning sessions. Monkeys were trained on the behavioral task with different images that were never used during scanning.

Prior to each scanning session, monkeys were intravenously injected with a contrast agent: monocrystalline iron oxide nanoparticle (MION, Feraheme (ferumoxytol), AMAG Pharmaceuticals, Inc., Waltham, MA, 30 mg per mL or BioPal Molday ION, Biophysics Assay Lab Inc., Worcester, MA, 30 mg per mL). MION to improves the contrast-to-noise ratio ∼3-fold (Vanduffel et al., 2001; Leite et al., 2002) and enhances spatial selectivity of MR signal changes (Zhao et al., 2006). MION was injected, approximately 30-60 min before scanning, into the saphenous vein below the knee (7 mg/kg), then flushed with a volume of sterile saline approximately double the volume of the MION injected. No additional MION was added during scanning, as MION has a long blood half-life (15.3 +/-3.5 hr) (Leite et al., 2002).

A Siemens 3T PRISMA MRI system with a custom six-channel surface coil (ScanMed, Omaha, NE) at the Brown University MRI Research Facility was used for whole-brain imaging. Anatomical scans consisted of a T1-MPRAGE (repetition time, TR, 2700 ms; echo time, TE, 3.16 ms; flip angle, 9°; 208 sagittal slices; 0.5 × 0.5 × 0.5 mm), a T2 anatomical (TR, 3200 ms; TE 410 ms; variable flip angle; 192 interleaved transversal slices; 0.4 × 0.4 × 0.4 mm), and an additional high resolution T2 anatomical (TR, 8020 ms; TE 44 ms; flip angle, 122°; 30 interleaved transversal slices; 0.4 × 0.4 × 1.2 mm). Functional images were acquired using a fat-saturated gradient-echoplanar sequence (TR, 1.8 s; TE, 15 ms; flip angle, 80°; 40 interleaved axial slices; 1.1 × 1.1 × 1.1 mm).

The target sample size (number of runs per monkey) was calculated using pilot data from a previous version of this task not included in the current data set. A region of interest was constructed from a cluster of deviant > NISR activation and the number of runs calculated for a significant effect in this region (using the beta values of the onset GLM, see below) at 80% power and alpha = 0.05 (G-Power). Guided by this power analysis and similar studies (Wang et al., 2015), we estimated a total of 200 runs across the three animals would be necessary.

### FMRI Data Analysis

The majority of the following analyses were performed in Matlab using SPM 12 (http://www.fil.Ion.ucl.ac.uk/spm). Prior to analysis, data were preprocessed using the following steps: reorienting (to ensure proper assignment of the x,y,z planes), motion correction (realignment), normalization, and spatial smoothing (2 mm isotropic Gaussian kernel separately for gray matter and white matter). All steps were performed on individual runs separately. The T1-MPRAGE anatomical image was skull stripped using FSL BET brain extraction tool (http://www.fmrib.ox.ac.uk/fsl/) to facilitate normalization. All images were normalized to the 112-RM SL macaque atlas (McLaren et al., 2009).

Runs were included for analysis only if they met the following criteria: the monkey had to be performing well and a sufficient number of acquisition volumes within the run had to pass data quality checks. The monkey’s performance was evaluated by calculating the percentage of time within a run that fixation was maintained. Runs were excluded if the monkey was fixating < 80% of the time (similar criteria as in (Vanduffel et al., 2001; Leite et al., 2002; Wang et al., 2015). Approximately 20% of runs were excluded due to poor fixation: 10% from monkey J, 3% from monkey W and 7% from monkey B. To evaluate data quality, we used the ART toolbox (Artifact Detection Tools, https://www.nitrc.org/projects/artifact_detect) to detect outlier volumes. Any volumes that had motion greater than one voxel (1.1 mm) in any direction were excluded. Any run with greater than 12% of volumes excluded was excluded from analysis (0% runs excluded for monkey J, 0.5% of runs excluded for monkey W, and 15% of runs excluded for monkey B). Runs with poor image quality due to artifact or banding to pre-process or analyze were also excluded. These accounted for 2% of the data for monkey J, 5% for monkey W, and 0.5% for monkey B. After applying these criteria, a total of 232 runs (average of 340 volumes per run for all animals, 93 sessions in total across animals) were included for analysis from 3 monkeys: monkey W: 97 runs (32 sessions); monkey J: 65 runs (32 sessions), and monkey B: 70 runs (32 sessions).

#### Models

Within-subject statistical models were constructed under the assumptions of the general linear model (GLM) in SPM 12 for each pseudo-subject bin. For all models, data were binned into approximately 10-run pseudo-subject bins. Each bin contained data from only one monkey. Runs were pseudo-randomly assigned to bins to balance the number of runs which followed each of the two sequential rules (three same one different or four same) and the distribution of runs from earlier and later scanning sessions. Condition regressors were all convolved with a gamma function (shape parameter = 1.55, scale parameter = 0.022727) to model the MION hemodynamic response function (Vanduffel and Farivar, 2014). The first six sequences in a run and reward times were included as nuisance conditions. Additional nuisance regressors were included for the six motion estimate parameters (translation and rotation), outlier volumes, and image variability (standard deviation of within run image movement variability, calculated using the ART toolbox). Outlier volumes were determined using the ART toolbox (standard global mean; global signal detection outlier detection threshold = 4.5; motion threshold = 1.1mm; scan to scan motion and global signal change for outlier detection) and one additional regressor with a “1” at only that volume was included for each volume to be “scrubbed”.

Regressors were estimated using a bin-specific fixed-effects model. Whole-brain estimates of bin-specific effects were entered into second-level analyses that treated bin as a random effect. One-sample t-tests (contrast value vs zero, p < 0.005) were used to assess significance. These effects were corrected for multiple comparisons when examining whole-brain group voxelwise effects using extent thresholds at the cluster level to yield false discovery rate (FDR) error correction (p < 0.05). Group contrasts were rendered on an inflated MNI canonical brain using Caret (Van Essen et al., 2001). Prior to selecting GLM’s we used the model assessment, comparison, and selection toolbox (MACS, https://github.com/JoramSoch/MACS, (Soch and Allefeld, 2018) to determine models that would be the best fit. Three GLMs were applied to the data as follows:

##### Onsets Model

To assess the univariate effects of deviant sequences, we constructed a model using instantaneous stimulus onset regressors for the first item in each sequence with the following nine condition regressors for different sequence types: short, medium, and long habituation sequence timing templates; NISR; rule deviants; two- and six-item number deviants; and two- and six-item rule and number deviants.

##### Parametric Last Item versus Unique Ramp Model

To directly test whether variance could be better accounted for by a phasic response at the last item in the sequence or ramping activation, we constructed a pair of models to allow last item and ramp regressors to compete for variance within the same model. Onset regressors were constructed with an instantaneous stimulus onset regressor at each position in the sequence with the same nine condition regressors for the different sequence types as in the Onsets Model: short, medium, and long habituation sequence timing templates; NISR; rule deviants; two- and six-item number deviants; and two- and six-item rule and number deviants. Including an onset at each position effectively modeled sustained activation throughout the sequence and enabled the inclusion of the following parametric regressors. The last item parametric was added as ones at the first sequence positions and an arbitrarily larger value (6) at the last item. The ramp parametric was entered as the sequence position (1-4, 1-2, or 1-6) for each sequence. Parametric regressors were implemented hierarchically in the GLM. Therefore, variance explained by the last parametric regressor (in this case, ramping), is above and beyond what could be explained by the onsets or last item regressors.

##### Parametric Ramp versus Unique Last Item Model

This second model of the pair sought to identify variance uniquely explained by the last item regressor, above and beyond variance explained by the onsets or ramping regressors. All other aspects of the model were the same as the unique ramp model above.

#### ROI Analyses

The primary bilateral regions of interest were constructed from the coordinates of a seed region centered in macaque monkey area 46d. These coordinates were determined, using diffusion weighted and functional MRI, to be most similar to the lateral portion of human area 10 (Gilbert et al., 2010; Sallet et al., 2013). Human lateral area 10 overlaps with areas of ramping activation observed in human RLPFC in previous studies (Desrochers et al., 2015, 2019; McKim and Desrochers, 2022). A 5 mm sphere was created around the center coordinate for the seed region in macaque Montreal Neurological Institute (MNI) space. The sphere was then transformed into 112RM-SL space using RheMap (Sirmpilatze & Klink, 2020, resulting in a sphere centered at *xyz* = 12.7, 32.6, 22.5 in 112RM-SL space. For identification of brain areas we also utilized the NIMH Macaque Template (NMT v02, Macaque Atlas, Jung et al., 2021; Seidlitz et al., 2018).

Additional ROIs were constructed with the explicit purpose of comparing nearby regions in DLPFC that were significant clusters of activation last item versus ramping models. Specifically, the significant left DLPFC cluster of activation for Unique Ramp, Rule Deviants > NISR in the unique ramp model (center *xyz* = −12.2, 36, 23) and the significant left DLPFC cluster of activation for Unique Last, Rule Deviants > NISR in the unique last item model (center *xyz* = − 12.3, 42.9, 21.8) were taken for comparison.

To compare activation within and across ROIs in a manner that controlled for variance, we extracted t-values from the condition of interest over baseline using the Marsbar toolbox (Jean-Baptiste Poline, 2002). T-values (one for each pseudo-subject bin, n = 22 bins) were entered into RM-ANOVAs with the identity of the monkey entered as a covariate.

## Results

Three monkeys (*macaca mulatta*) performed no-report abstract sequence viewing while undergoing awake fMRI scanning. The monkeys were trained to fixate on a central spot while viewing a stream of fractal images arranged into four-item visual sequences (based on Wang, et al., **Figure 1**). This task did not require responses, only fixation, and thus was termed “no-report”. The task was performed in runs (∼10 min each), that each contained five blocks. For each run, the first block habituated animals to one of two possible sequential rules AAAB, or AAAA (A and B represent different images drawn from a pool of four possible images; 30 sequences in total per block). Habituation sequences each had one of six possible timing templates to balance stimulus and sequence durations across sequence types. Each subsequent block contained rare deviants (6 of the 30 sequence repetitions per block) of one of the following four possible types: new images following the same rule, number deviants (2 or 6 items), rule deviants (e.g., AAAA), or double deviants. All deviant images were drawn from a separate three-image pool. The five total blocks were interleaved with 16 s fixation blocks. To encourage animals to maintain fixation throughout, reward was administered on a graduated schedule not correlated with sequence presentation: the longer they maintained fixation, the shorter the duration between rewards. Reward was thus decorrelated from the four-item visual sequences. A total of 232 runs were analyzed (97 monkey W, 65 monkey J, 70 monkey B). Monkeys performed the task well and fixated for 95% of the time in included runs (see Methods for those excluded).

### Monkey DLPFC represents changes in abstract visual sequences

Our first goal was to test whether area 46 differentially responds when there is a change in the abstract visual sequence. Because this task is no-report, we examined this question using neural responses (BOLD) to deviant sequences. Previous work has shown that such deviant responses disappear with inattention and are robust in brain areas processing sequence related information (Bekinschtein et al., 2009; Dehaene et al., 2015). These results suggest that deviant neural responses indicate that individuals are attending to the sequences, even in the absence of a report. Therefore, we used neural responses to rare deviant sequences to indicate awareness of changes to an established abstract sequence, as is in similar auditory tasks (Uhrig et al., 2014; Wang et al., 2015). To specifically query the responses to abstract sequence changes, we could not simply compare habituated sequences to deviant sequences, as the deviant sequences were composed from a different pool of images than the habituation sequences, and any differences observed between habituation and deviant sequences could have resulted from differences in image identity. Therefore, to specifically examine changes in abstract sequence structure, we compared new items of the same rule (NISR) to number deviants and rule deviants. All images in this comparison were drawn from the same pool of (deviant) images. Double deviants were not included in analyses because of the inability to dissociate between changes due to rule and number.

We first constructed an unbiased region of interest (ROI) for monkey area 46 in each hemisphere to compare activity between rule and number deviants and NISR. Monkey area 46 has many potential functional subdivisions (Gerbella et al., 2010, 2013; Borra et al., 2011, 2019; Saleem et al., 2014); therefore, we created a 5 mm sphere centered on a seed region identified as having the most similar connectivity with human RLPFC in monkey diffusion and functional MRI (Sallet et al., 2013, center *xyz* = 12.7, 32.6, 22.5 in area 46d, see Methods). The resulting sphere spanned a small region of area 46 that encompassed 46d, 46f, and 46v (NIMH Macaque Template, NMT v2.0 Macaque Atlas, Jung et al., 2021; Seidlitz et al., 2018). Because sequence related activity in human RLPFC was observed in both hemispheres, we used identical spheres (mirrored coordinates) in the left and right hemispheres (referred to as L46 and R46, respectively) throughout. To compare activity between rule and number deviants and NISR in these ROIs, we created a model that included separate regressors for each habituation timing and deviant type, modeled as zero-duration onsets. Statistical testing was performed on ∼10 run bins (n = 22), each consisting of data from a single monkey (see Methods). We compared t-values from the contrast of each condition over baseline (e.g., Rule Deviants > Baseline vs. NISR > Baseline) to account for potential differences in variance across conditions. This type of comparison was used to examine ROI activity throughout, and we refer to comparisons by the conditions of interest (without listing the contrast over baseline, e.g., Rule Deviants > NISR). All statistical tests on ROIs were performed on binned data and included a covariate for monkey identity (n = 3). While we report the effect of monkey in the following analyses, the main focus of the study was not on individual differences, and our discussion focuses on condition effects.

We found that R46 represented abstract sequence changes, showing greater deviant activation across both deviant types (**Figure 2**;**Table 1**). Responses were reliably greater for rule deviants compared to NISR (sequence type: F(1, 19) = 4.6, p = 0.046, η_p_^2^ = 0.19) and marginally greater for number deviants compared to NISR (sequence type: F(1, 19) = 3.9, *p* = 0.062, η_p_^2^ = 0.17). Even though deviant responses compared to NISR in L46 did not reach statistical significance (**Table 1**), there were no reliable differences between responses in R46 and L46 (**Table 2**). These results suggest that a specific region of monkey DLPFC, area 46, monitors abstract visual sequence structure.

**Table 1.**
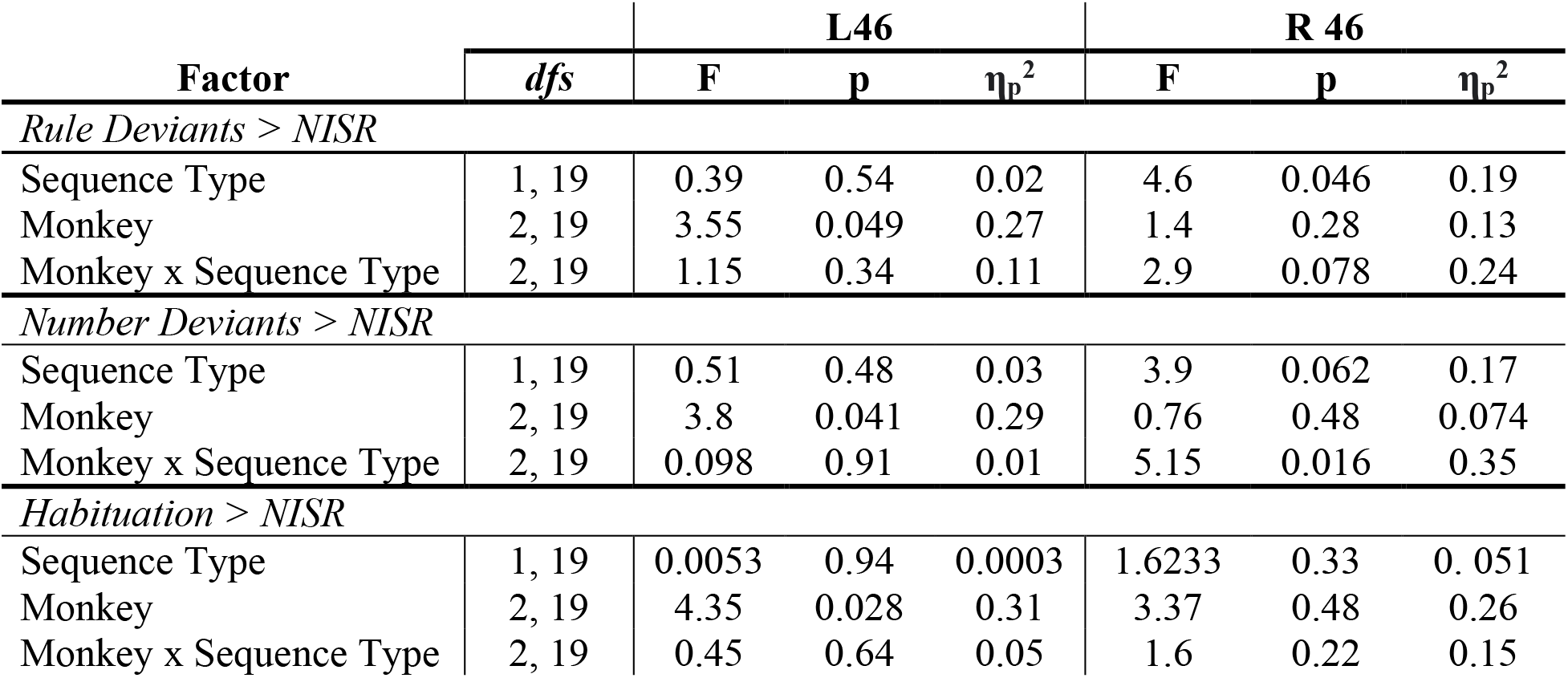
Repeated measures ANOVAs comparing rule and number deviants to NISR in L46 and R46.

**Table 2.**
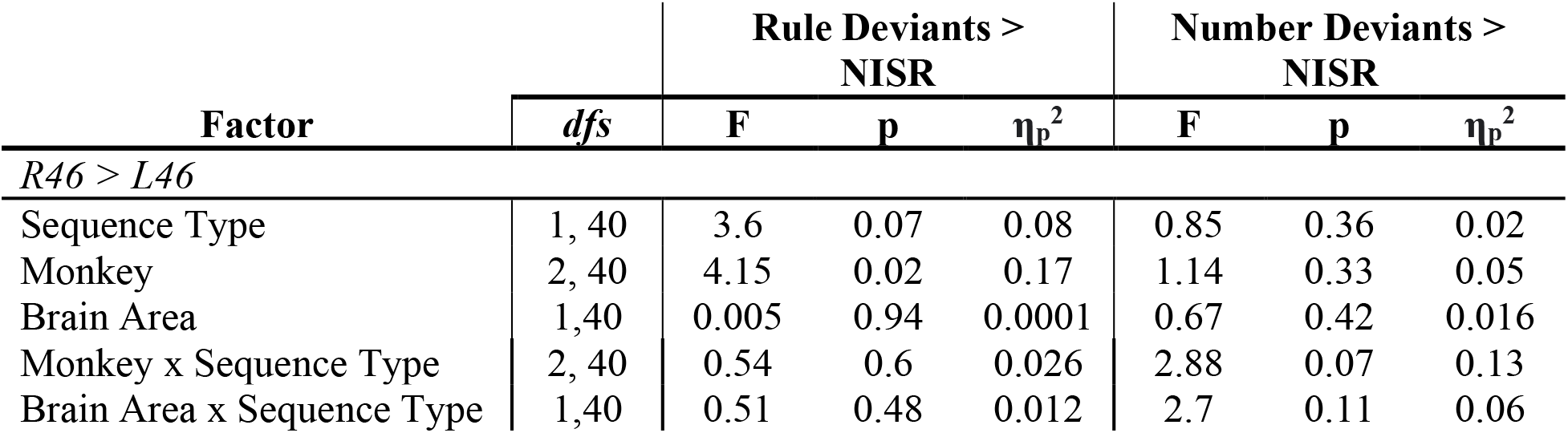
Repeated measures ANOVAs comparing deviant responses in L46 and R46.

|  |  | Rule Deviants ><br>NISR |  |  | Number Deviants ><br>NISR |  |  |
| --- | --- | --- | --- | --- | --- | --- | --- |
| Factor | dfs | F | p | $\eta_p^2$ | F | p | $\eta_p^2$ |
| *R46 > L46* |  |  |  |  |  |  |  |
| Sequence Type | 1, 40 | 3.6 | 0.07 | 0.08 | 0.85 | 0.36 | 0.02 |
| Monkey | 2, 40 | 4.15 | 0.02 | 0.17 | 1.14 | 0.33 | 0.05 |
| Brain Area | 1,40 | 0.005 | 0.94 | 0.0001 | 0.67 | 0.42 | 0.016 |
| Monkey x Sequence Type | 2, 40 | 0.54 | 0.6 | 0.026 | 2.88 | 0.07 | 0.13 |
| Brain Area x Sequence Type | 1,40 | 0.51 | 0.48 | 0.012 | 2.7 | 0.11 | 0.06 |

**Figure 2.**
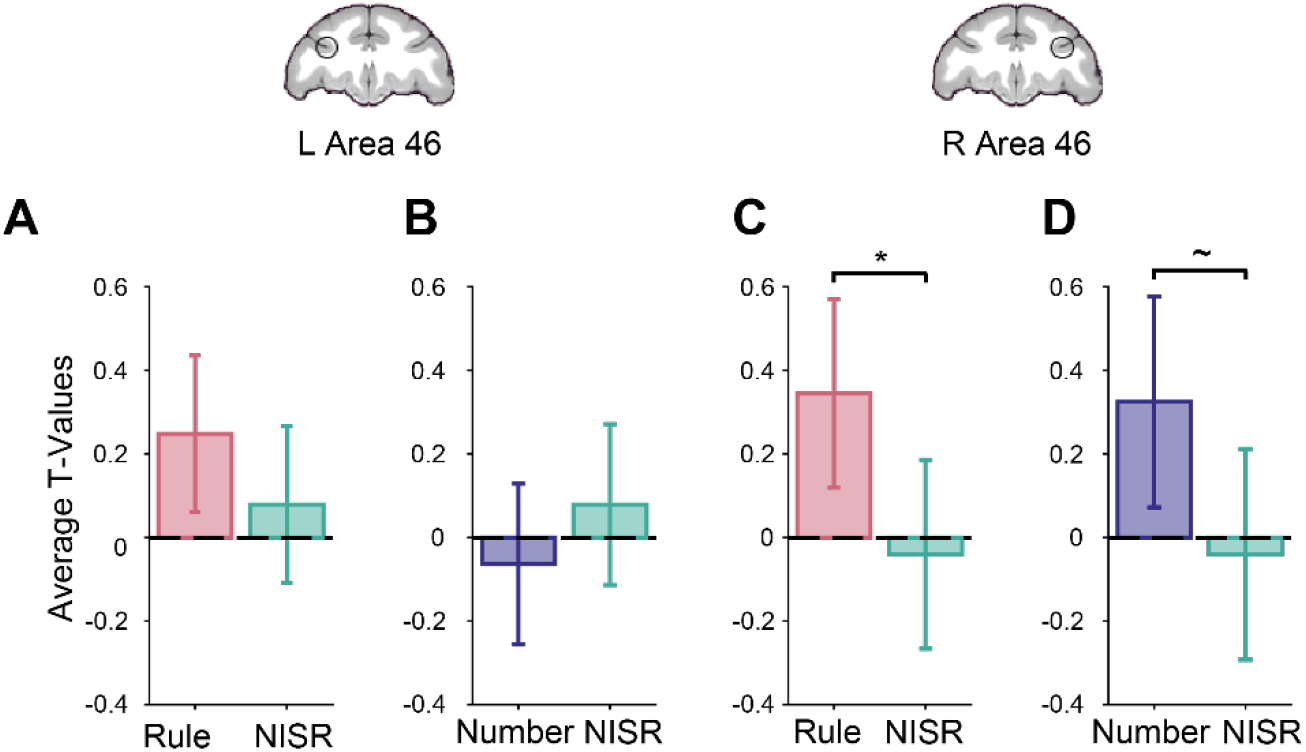
Area 46 represents abstract visual sequences. T-values for the condition of interest > baseline are shown. The locations of area 46 regions of interest (ROIs), L46 and R46, are outlined in black on coronal sections (y = 33). **A**. Rule deviants compared to new items, same rule (NISR) in L46. **B**. Number deviants compared to NISR in L46. **C**. Rule deviants compared to NISR in R46 showed a reliable difference. **D**. Number deviants compared to NISR in R46 showed a marginal difference. Comparisons in L46 showed similar trends as in R46. Error bars are 95% confidence intervals (1.96 x standard error of the within-bin mean).

As a control, we also examined conditions where the pool of images differed, but the abstract sequential structure did not. If area 46 was responding specifically to a change in the abstract sequential structure, then a change in the images should not change its activation level. We examined the difference in contrast t-values between NISR and habituation trials with comparable stimulus durations (“medium” timing, as in **Figure 1D**). We did not find any significant differences between these conditions in either R46 or L46 (**Table 1**), indicating that changes in activation in area 46 were specific to changes in abstract sequential structure.

Results from area 46 ROIs were supported by whole-brain contrasts examining responses to number and rule deviants. Contrasts of Rule Deviants > NISR and Number Deviants > NISR both showed significant clusters of activation in right area 46 (**Figure 3, Table 3, Extended Data Figure 3-1**). Other significant clusters of activation were located in areas such as the caudate nucleus, high-level auditory cortex (rostromedial belt region), and dorsal premotor cortex, areas also observed in a similar auditory sequence task (Wang et al., 2015). Further, deviant responses in earlier sensory areas (e.g., V2) may be analogous to responses in auditory cortex previously observed. Though we could not address the question of sensory generality within the current experiment, these results raise the intriguing possibility that these previously indicated areas could be sensory-modality general in their responses to abstract sequential structure.

**Table 3.**
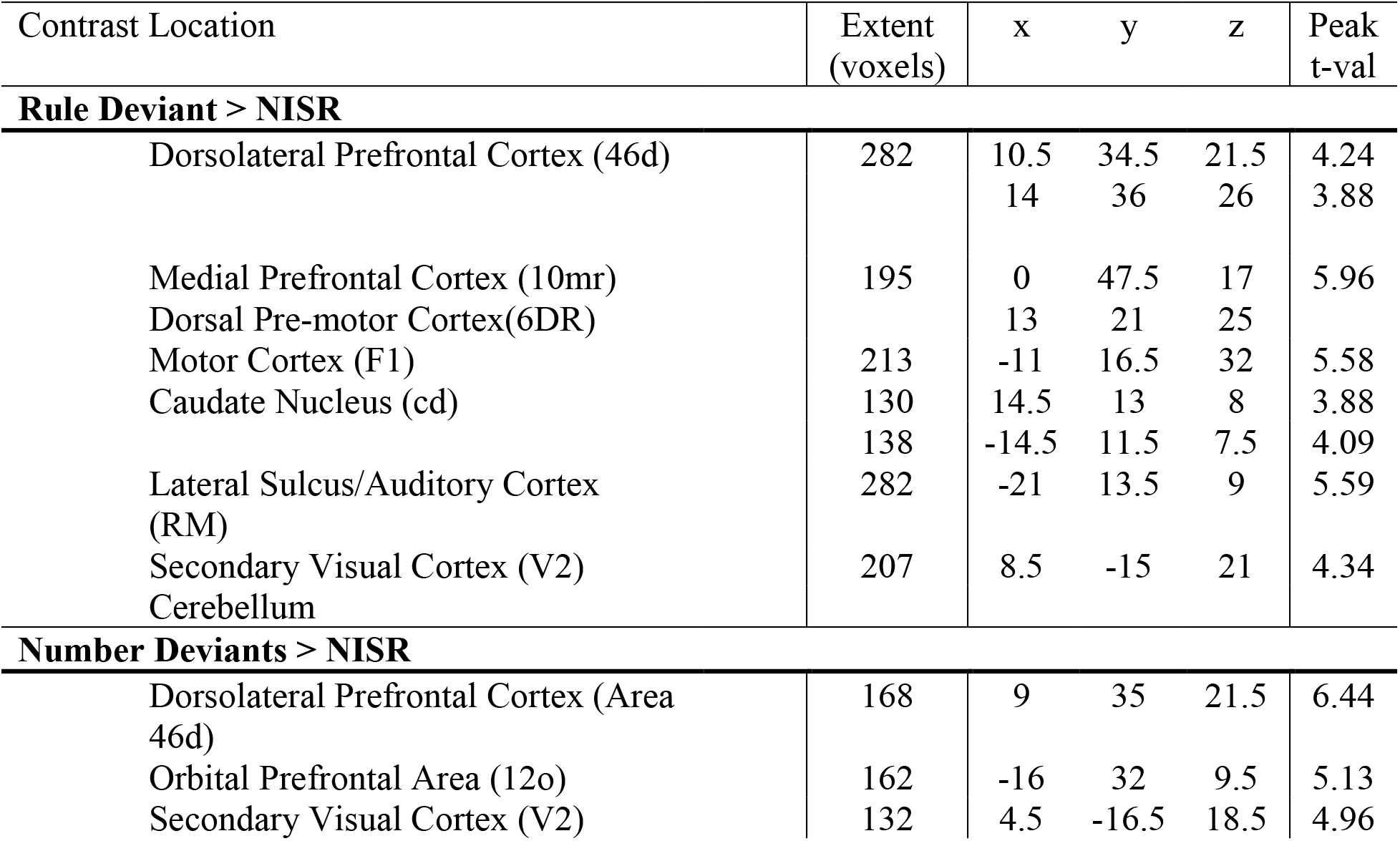
Rule and number deviants compared to NISR contrast activation coordinates. Area labels as in the NIMH NMT v02 Macaque Atlas.

**Figure 3.**
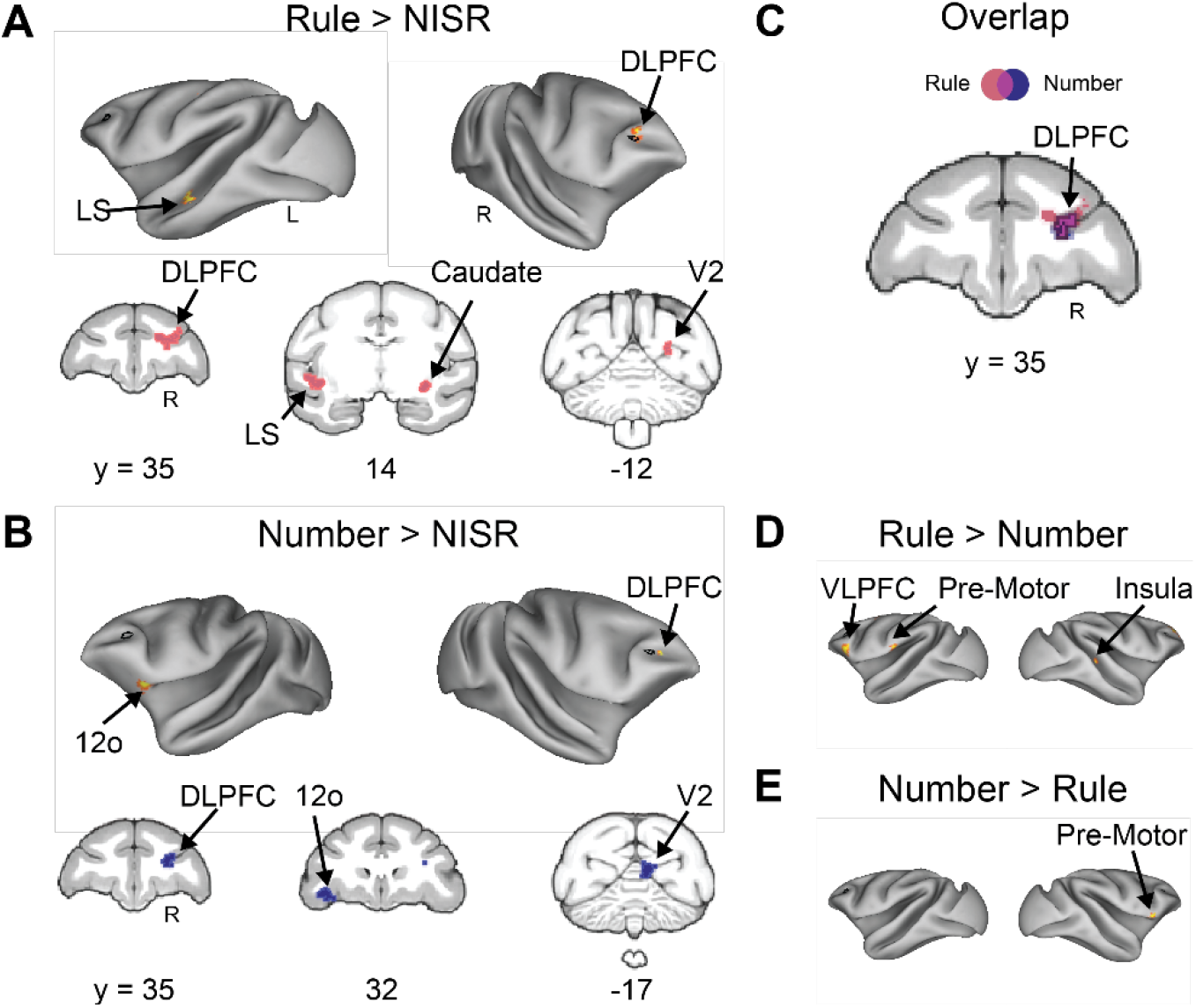
Whole-brain deviant activity shows area 46 represents both rule and number deviants. **A**. Voxel wise contrast of Rule Deviants > NISR false discovery rate (FDR) error cluster corrected for multiple comparisons (FDRc < 0.05, height p < 0.005 unc., extent = 130) are shown. Individual monkey data shown in **Extended Data Figure 3-1A. B**. Voxelwise contrast of Number Deviants > NISR (FDRc < 0.05, height p < 0.005 unc., extent = 132). Individual monkey data shown in **Extended Data Figure 3-1B. C**. Overlap of Rule Deviants > NISR and Number Deviants > NISR contrasts showed significant, unique conjunction (violet outlined in black) only in the DLPFC. **D**. Voxelwise contrast of Rule Deviants > Number Deviants showing no significant clusters of activation in area 46 (FDRc < 0.05, height p < 0.005 unc., extent = 102). **E**. As in (C) for Number Deviants > Rule Deviants (FDRc < 0.05, height p < 0.005 unc., extent = 111). Black outline on inflated brains indicates location of L46 or R46 (depending on the hemisphere shown) for reference. Lateral Sulcus (LS), Dorsal Lateral Pre-frontal Cortex (DLPFC), Ventral Lateral Pre-frontal Cortex (VLPFC), Secondary Visual Cortex (V2), Orbital Prefrontal Area (12o).

To determine if the observed responses to number and rule deviants were similar in area 46, we directly examined whether these responses generalized across deviant types. The t-values in R46 were not different between rule and number deviants (sequence type: F(1,19) = 0.0011, *p* = 0.92, η_p_^2^ = 0.11, monkey: F(2, 19) = 2.5, p = 0.11, η_p_^2^ = 0.21; monkey x sequence type: F(2, 19) = 1.15, p = 0.34, η_p_^2^ = 0.11). Next, we performed a conjunction analysis to determine the areas of activation that overlapped in the Rule Deviants > NISR and Number Deviants > NISR contrasts. We found that the only cluster of significant overlap between the deviant contrasts was in right area 46 (**Figure 3C**). In support of this finding, whole-brain direct contrasts of rule and number deviants showed significant activation clusters in VLPFC, insula, and pre-motor cortex, but no significant clusters in area 46 (**Figure 3D, E**). In summary, these results suggest that abstract visual sequential structure is monitored in area 46, and that this monitoring is both unique to area 46 and general across different kinds of deviations.

### Ramping activation reflects sequence monitoring in monkey DLPFC

We next tested the prediction that area 46 would display similar dynamics to those observed in humans during abstract sequences. Specifically, previous experiments in humans showed that BOLD activity increased (“ramped”) from the beginning to the end of sequences in the RLPFC (Desrochers et al., 2015, 2019; McKim and Desrochers, 2022). Given the similarity in connectivity between monkey area 46 and human RLPFC, we hypothesized that changes in abstract sequence structure would also produce changes in ramping activation in area 46 if abstract sequence monitoring underlies this dynamic.

To test if ramping dynamics were present in area 46 during this task, we first designed a model to isolate these dynamics. This model included regressors for the three dominant potential dynamics (**Figure 4A, Extended Data Figure 4-1**, see also Materials and Methods). First, instantaneous onsets were included for each image presentation, effectively modeling sustained activation throughout each sequence. Then, two parametric regressors were included: last item change and ramping. The ramping regressor parallels the analysis that revealed ramping in human RLPFC: it increases linearly from the first to the last item in the sequence, and resets at each new sequence. The last item change regressor is low at the first three positions of each sequence, and high at the last item in the sequence. This regressor was designed to account for the fact that differences in the rule that the sequence followed would occur at the last item (e.g., the difference between AAAA and AAAB occurs at the fourth item), and a dynamic associated with this change could have variance mistakenly assigned to a ramping regressor.

**Figure 4.**
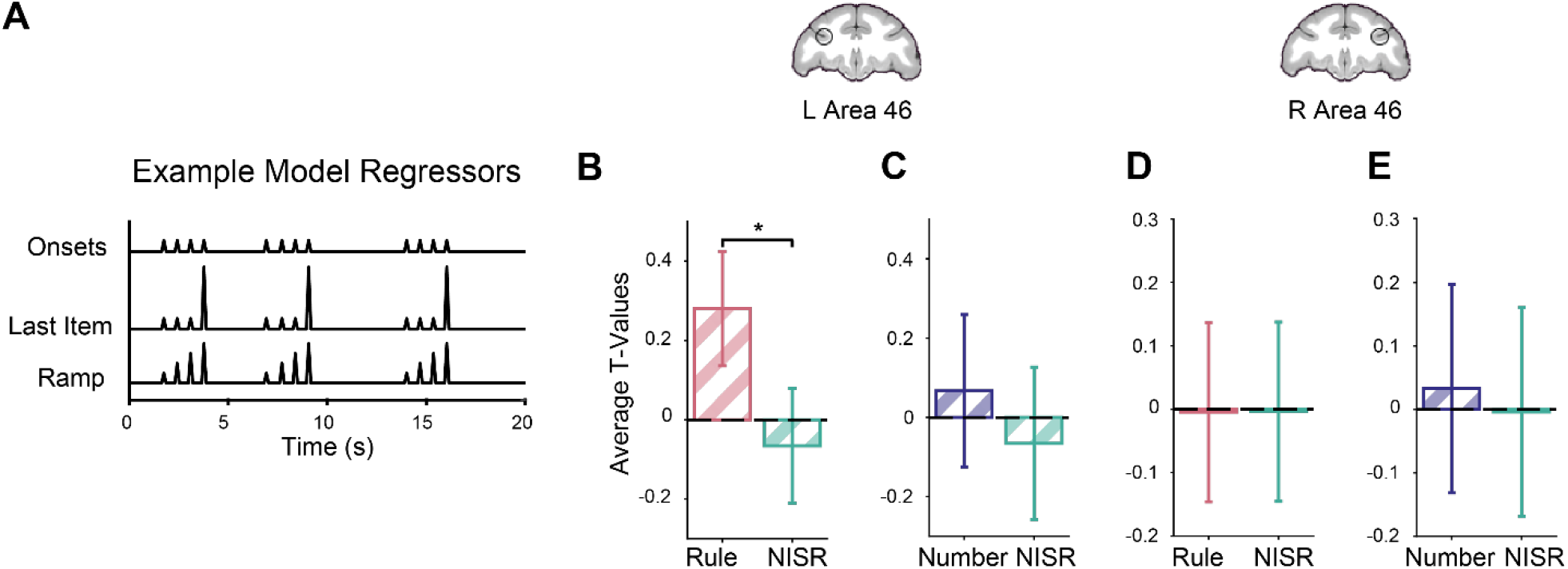
Area 46 shows ramping activity for deviations to an established sequence rule. Parametric models and T-values for the condition of interest > baseline shown. Coronal brain slices (*y* = 33) show locations of area 46 ROIs, L46 and R46, outlined in black. **A**. Example of regressors used to model parametric ramp and parametric last item. Example regressors through the orthogonalization process shown in **Extended Data Figure 4-1. B**. Unique ramping during rule deviants compared to NISR in L46 showed a reliable difference. **C**. Unique ramping during number deviants compared to NISR in L46. **D**. Unique ramp number deviants compared to NISR in R46. **E**. Unique ramping during number deviants compared to NISR in R46. Comparisons that were not reliably different showed similar trends. Error bars are 95% confidence intervals (1.96 x standard error of the within-bin mean).

These parametric regressors were orthogonalized in a stepwise fashion and only absorbed variance above and beyond variance accounted for by the onset regressor. The last regressor, therefore, contained “unique” variance. A pair of models, one with unique variance assigned to ramp and one with it assigned to last item, were created to examine these dynamics. The correlation between the resulting regressors was, as expected, low. For example, in one bin of this parametric model with the unique variance assigned to ramp, the average correlation coefficient between last item change and ramp regressors was 0.00005 (± 0.0001 standard deviation). While there are likely nearly infinite variations of dynamics possible that lie across a spectrum between the last item change and ramping (e.g., exponential), our purpose in designing these models was not to explore the space of all possible dynamics, but to test for ramping dynamics in area 46.

We found ramping dynamics in monkey area 46 related to abstract sequence monitoring. When comparing t-values of contrasts between deviants and baseline using the same spherical area 46 ROIs described above, we observed significant, unique variance ascribed to ramping activation in L46 during rule deviants compared to NISR (sequence type: F(1, 19) = 5.03, *p* = 0.037, η_p_^2^ = 0.21; **Table 4**; **Figure 4B**). Unique ramping activation showed a numerical trend in the same direction for number deviants compared to NISR in L46, but it did not reach statistical significance (**Table 4**). Activity in L46 was not reliably different between the two deviant types (sequence type: F(1,19) = 2.06, *p* = 0.17, η_p_^2^ = 0.098, monkey: F(2, 19) = 0.57, p = 0.64, η_p_^2^ = 0.06; monkey x sequence type: F(2, 19) = 2.01, p = 0. 16, η_p_^2^ = 0.17). In R46, changes in unique ramping activation during rule and number deviants were not significant (**Table 4**). Despite apparent differences between L46 and R46, unique ramping was not reliably different between these ROIs (**Table 5**). These results suggest that area 46 shows ramping dynamics for sequential rule changes. Interestingly, ramping may be preferentially present in L46, suggesting that while both hemispheres detect abstract sequence deviations, they may do so with different dynamics. Further, these results suggest similar sequential monitoring processes may be present across species in analogous areas.

**Table 4.** Repeated measures ANOVAs comparing unique ramping activity during rule and number deviants to NISR in L46 and R46.

| Factor | dfs | L46 |  |  | R 46 |  |  |
| --- | --- | --- | --- | --- | --- | --- | --- |
| | | F | p | $\eta_p^2$ | F | p | $\eta_p^2$ |

| *Unique Ramp, Rule Deviants > NISR* |  |  |  |  |  |  |  |
| --- | --- | --- | --- | --- | --- | --- | --- |
| Sequence Type | 1, 19 | 5.03 | 0.037 | 0.21 | 0.01 | 0.91 | 0.00075 |
| Monkey | 2, 19 | 0.82 | 0.45 | 0.08 | 0.64 | 0.54 | 0.63 |
| Monkey x Sequence Type | 2, 19 | 0.23 | 0.8 | 0.0235 | 0.81 | 0.46 | 0.08 |
| *Unique Ramp, Number Deviants > NISR* |  |  |  |  |  |  |  |
| Sequence Type | 1, 19 | 0.42 | 0.52 | 0.02 | 0.03 | 0.86 | 0.0016 |
| Monkey | 2, 19 | 2.39 | 0.12 | 0.2 | 0.24 | 0.45 | 0.08 |
| Monkey x Sequence Type | 2, 19 | 5.15 | 0.63 | 0.047 | 5.15 | 0.84 | 0.0235 |

**Table 5.** Repeated measures ANOVAs comparing unique ramping deviant responses in L46 and R46.

|  |  | Unique Ramp, Rule<br>Deviants > NISR |  |  | Unique Ramp, Number<br>Deviants > NISR |  |  |
| --- | --- | --- | --- | --- | --- | --- | --- |
| Factor | dfs | F | p | $\eta_p^2$ | F | p | $\eta_p^2$ |
| *R46 > L46* |  |  |  |  |  |  |  |
| Sequence Type | 1, 40 | 0.38 | 0.54 | 0.009 | 2.4 | 0.13 | 0.06 |
| Monkey | 2, 40 | 1.62 | 0.21 | 0.07 | 1.5 | 0.24 | 0.07 |
| Brain Area | 1, 40 | 0.02 | 0.89 | 0.0005 | 1.5 | 0.25 | 0.03 |
| Monkey x Sequence Type | 2, 40 | 0.64 | 0.53 | 0.03 | 0.12 | 0.89 | 0.006 |
| Brain Area x Sequence Type | 1, 40 | 0.14 | 0.71 | 0.003 | 2.7 | 0.11 | 0.06 |

Because variance due to changes at the last item of the sequence could be misattributed to ramping regressors, we directly compared activity in area 46 that could be accounted for by ramping and last item change regressors. In this control analysis, we found that activity was significantly greater for unique ramping than unique last item change during rule deviants in L46 (sequence type: F(1, 19) = 9.53, *p* = 0.006, η_p_^2^ = 0.33; **Table 6**; **Figure 5A**). Number deviants in L46 and both deviants in R46 showed similar numerical trends for unique ramping accounting for greater variance than last item change but did not reach statistical significance (**Figure 5B-D**; **Table 6**). These results suggest that area 46 dynamics during abstract sequence monitoring are best accounted for by a ramping function, rather than a last item change.

**Table 6.** Repeated measures ANOVAs comparing unique ramping activity to unique last item activity during rule and number deviants to NISR in L46 and R46.

| Factor | dfs | L46 |  |  | R 46 |  |  |
| --- | --- | --- | --- | --- | --- | --- | --- |
| | | F | p | $\eta_p^2$ | F | p | $\eta_p^2$ |
| *Unique Ramp, Rule Deviants > Unique Last Item, Rule Deviants* |  |  |  |  |  |  |  |
| Sequence Type | 1, 19 | 9.53 | 0.006 | 0.33 | 0.008 | 0.93 | 0.0004 |
| Monkey | 2, 19 | 1.02 | 0.38 | 0.97 | 0.72 | 0.5 | 0.07 |
| Monkey x Sequence Type | 2, 19 | 0.37 | 0.7 | 0.38 | 0.09 | 0.91 | 0.01 |
| *Unique Ramp, Number Deviants > Unique Last Item, Number Deviants* |  |  |  |  |  |  |  |
| Sequence Type | 1, 19 | 0.196 | 0.66 | 0.01 | 0.07 | 0.8 | 0.0036 |
| Monkey | 2, 19 | 2.08 | 0.15 | 0.18 | 1.47 | 0.25 | 0.13 |
| Monkey x Sequence Type | 2, 19 | 1.26 | 0.31 | 0.12 | 0.15 | 0.86 | 0.015 |

**Figure 5.**
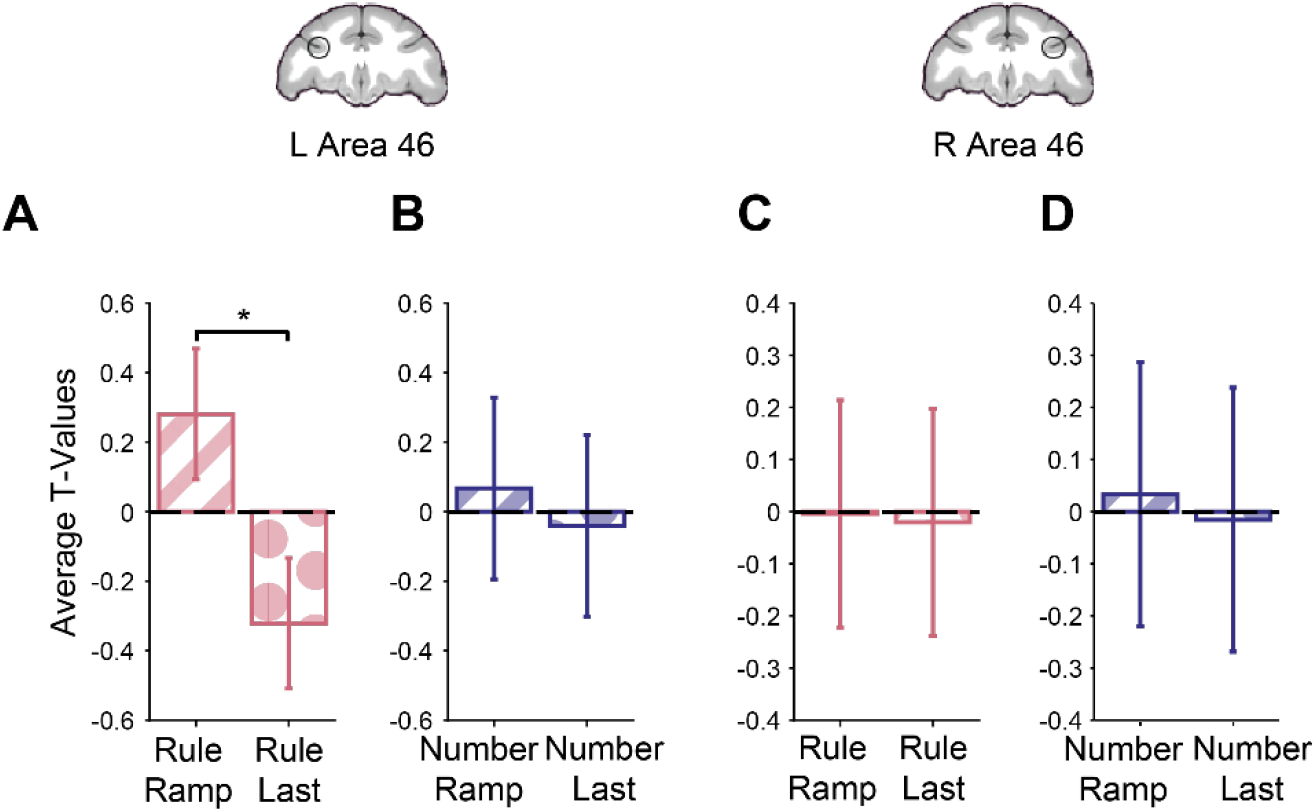
Area 46 shows greater unique ramping than unique last item activity during abstract sequence deviants. T-values for the condition of interest > baseline shown. **A**. Unique ramp compared to unique last item during rule deviants in L46 showed a reliable difference. **B**. Unique ramp compared to unique last item during number deviants in L46. **C**. Unique ramp compared to unique last item during rule deviants in R46. **D**. Unique ramp compared to unique last item during number deviants in R46. Comparisons that were not reliably different showed similar trends. Error bars are 95% confidence intervals (1.96 x standard error of the within-bin mean).

ROI results were supported by whole-brain contrasts that examined ramping and last item dynamics. Unique ramping was present in left area 46 during rule deviants compared to NISR (**Figure 6A, Table 7, Extended Data Figure 3-1C**). Other clusters of activation were present in the visual cortex and superior temporal gyrus that were similar to those observed for ramping activation in humans (Desrochers et al., 2015, 2019). As expected from the ROI analyses, the number deviant ramping contrast had significant whole brain clusters in areas such as visual cortex and putamen but no significant clusters in area 46 (**Figure 6B, Table 7**). The localization of significant unique last item clusters was different than ramping. Specifically, a more anterior region of ventral area 46 (46v), in contrast to more posterior and dorsal area 46 (46d) observed for unique ramping, showed significant activation for unique last item change in rule deviants compared to NISR (**Figure 7A, Table 8**). Other significant clusters of activation for last item change included the somatosensory cortex and central orbitofrontal cortex (area 13m). We also contrasted unique last item variance in number deviants compared to NISR, and observed activation in areas such as anterior cingulate gyrus, insula and visual cortices but no surviving clusters in area 46 (**Figure 7B, Table 8**). These results suggest that subregions of area 46 within the same hemisphere may reflect separable aspects of changes in sequential structure.

**Table 7.** Unique ramp rule and number deviants compared to NISR contrast activation coordinates.

| Contrast Location | Extent (voxels) | x | y | z | Peak t-val |
| --- | --- | --- | --- | --- | --- |
| **Unique Ramp, Rule Deviants > NISR** |  |  |  |  |  |
| Primary Motor Cortex (F1) | 153 | 11 | 13.5 | 29.5 | 6.48 |
| Caudal Dorsal Pre-Motor Cortex (PMdc) | 89 | 12 | 18 | 34.5 | 6.17 |
| Dorsal Dorsal Lateral Prefrontal Cortex (46d) | 130 | -12.5 | 36 | 23 | 5.34 |
| Intraparietal Area (IPa) | 121 | 16 | 18.5 | 2 | 5.33 |
| Rostral Area 12 (12r) | 128 | -15 | 29 | 14.5 | 5.19 |
| Caudate Nucleus (Cd) | 385 | -10.5 | 19 | 23 | 5.15 |
| Caudal Pre-Motor Cortex (PMdc) | 113 | -18.5 | 20 | 30 | 5 |
| Secondary Visual Cortex (V2) | 82 | 28 | -7.5 | 14.5 | 4.67 |
| Primary Visual Cortex (V1) | 101 | -20.5 | -11 | 27 | 4.66 |
| Body of the Fornix (bfX) | 120 | -1.5 | 12.5 | 22.5 | 4.49 |
| Somatosensory Parietal Cortex (1,2) | 120 | -13 | 7.5 | 33.5 | 4.4 |
| Primary Motor Cortex (M1) | 97 | -3.5 | 14.5 | 38 | 4.3 |
| Medial Septum (ms) | 82 | -1 | 18.5 | 13 | 4.18 |
| Secondary/Primary Visual Cortex (V2/V1) | 129 | 9 | -11 | 19 | 4.16 |
| Intraparietal Area (IPa) | 97 | 18.5 | 7.5 | 16 | 4.01 |
| Globus Pallidus (MGPi) | 250 | -6.5 | 13 | 8 | 3.99 |
| Pre-genua Cortex (24a) | 101 | -2 | 38.5 | 17 | 3.96 |
| **Unique Ramp, Number Deviants > NISR** |  |  |  |  |  |
| Visual Area 3 (V3) | 163 | 7.5 | -12.5 | 26.5 | 6.42 |
| Anterior Lateral Belt Region (AL) | 130 | 28 | 17 | 12.5 | 6.35 |
| Agranular and Dysgranular Insula (Ia/Id) | 103 | 21 | 19 | 10.5 | 5.62 |
| Putamen (Pu) | 127 | 7 | 24.5 | 10.5 | 5.5 |
| Caudate Nucleus (Cd) | 772 | -5.5 | 20 | 18 | 5.29 |
| Primary Visual Cortex (V1) | 132 | 9 | -11.5 | 19 | 5.16 |
| Cerebellum | 250 | 7 | -15.5 | 12 | 4.85 |
| Somatosensory Parietal Cortex (1,2) | 89 | -4.5 | 4 | 35 | 4.72 |
| Cerebellum | 138 | 4 | -18 | 5.5 | 4.67 |
| Anterior Intraparietal Area (AIP) | 123 | 18.5 | 10.5 | 25.5 | 4.65 |
| Somatosensory Parietal Cortex (1,2) | 115 | -21 | 13 | 32.5 | 4.5 |
| Caudal Dorsal Pre-Motor Cortex (PMdc) | 324 | -10.5 | 20 | 28 | 4.45 |
| Ventral Tegmental Area (VTA) | 282 | -4 | 4 | 3 | 4.37 |
| Posterior Cingulate Gyrus (23c) | 100 | -7.5 | 5.5 | 30.5 | 4.07 |
| Somatosensory Parietal Cortex (1,2) | 152 | -12.5 | 6.5 | 35 | 4.01 |

**Table 8.** Unique last item rule and number deviants compared to NISR contrast activation coordinates.

| Contrast Location | Extent<br>(voxels) | x | y | z | Peak<br>t-val |
| --- | --- | --- | --- | --- | --- |
| **Unique Last Item, Rule Deviants > NISR** |  |  |  |  |  |
| Ventral Dorsal Lateral Prefrontal Cortex (46v) | 148 | -12 | 43 | 22 | 5.23 |
| Granular Insula (Ig) | 170 | 19 | 11 | 20 | 5.06 |
| Medial Area 11 (11m) | 156 | 3.5 | 41 | 17.5 | 4.93 |
| Ventral Intraparietal Area/White Matter (VIP/WM) | 360 | 8.5 | -1 | 22.5 | 4.1 |
| **Unique Last Item, Number Deviants > NISR** |  |  |  |  |  |
| Area TFO (TFO) | 121 | -14 | 4 | 1.5 | 5.75 |
| Caudate Nucleus | 82 | 1 | 29.5 | 16.5 | 5.32 |
| Primary Visual Cortex (V1) | 247 | -8.5 | -18.5 | 12 | 5.13 |
| Area 13 (13m) | 184 | 4 | 38 | 18.5 | 4.92 |
| Cerebellum | 90 | 12.5 | -12.5 | 5 | 4.89 |
| Granular Insula (Ig) | 117 | 17.5 | 16 | 15.5 | 4.36 |
| Temporal Parietooccipital Associated Area (TPO) | 88 | 21.5 | 14 | 7.5 | 4.32 |
| Caudomedial Belt Region (CM) | 116 | -15 | 2.5 | 22 | 4.19 |
| Anterior Cingulate Gyrus (24a) | 85 | -2 | 28.5 | 20.5 | 4.14 |
| Posterior Cingulate Gyrus (29) | 151 | 7 | 0.5 | 20.5 | 4.03 |

**Figure 6.**
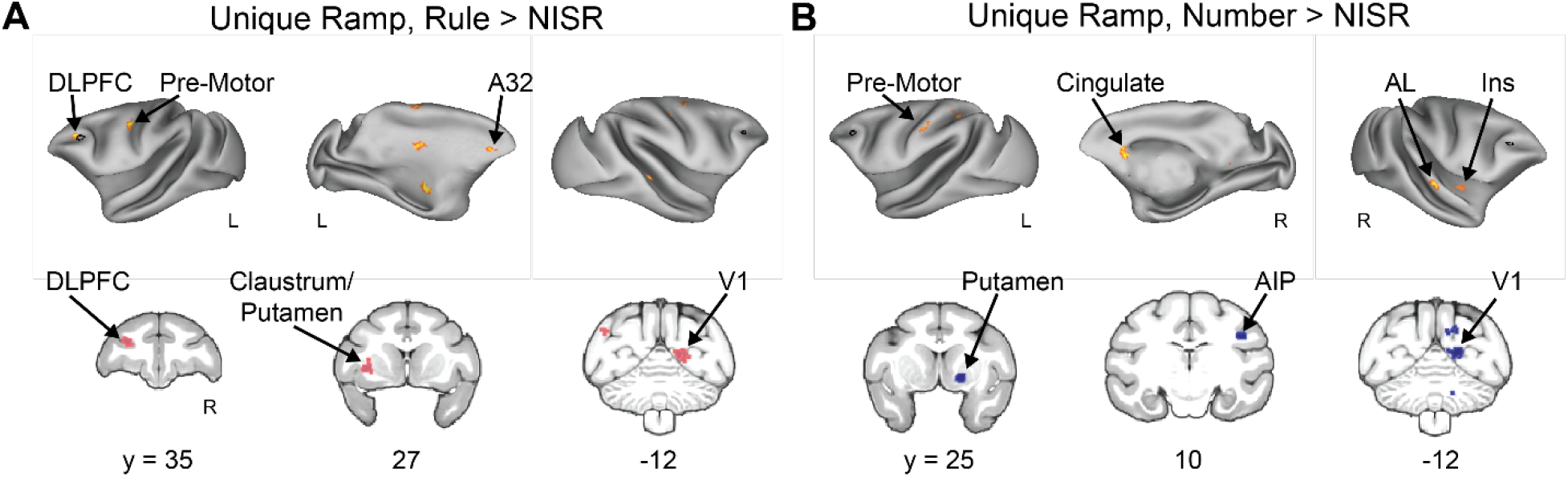
Area 46 shows unique ramping during abstract sequence deviants. **A**. Unique Ramp, Rule Deviants > NISR (FDRc p < 0.05, height p < 0.005 unc., ext. 82). Individual monkey data shown in **Extended Data Figure 3-1C. B**. Unique Ramp, Number Deviants > NISR (FDRc p < 0.05, height p < 0.005 unc., ext. 89). Black outline on inflated brains indicates location of L46 or R46 (depending on the hemisphere shown) for reference. Medial Prefrontal Cortex (A32) Dorsal Lateral Pre-frontal Cortex (DLPFC), Primary Visual Cortex (V1), Insular Cortex (Ins), Anterior Lateral Belt Region of the Auditory Cortex (AL), Anterior Intraparietal Area (AIP).

**Figure 7.**
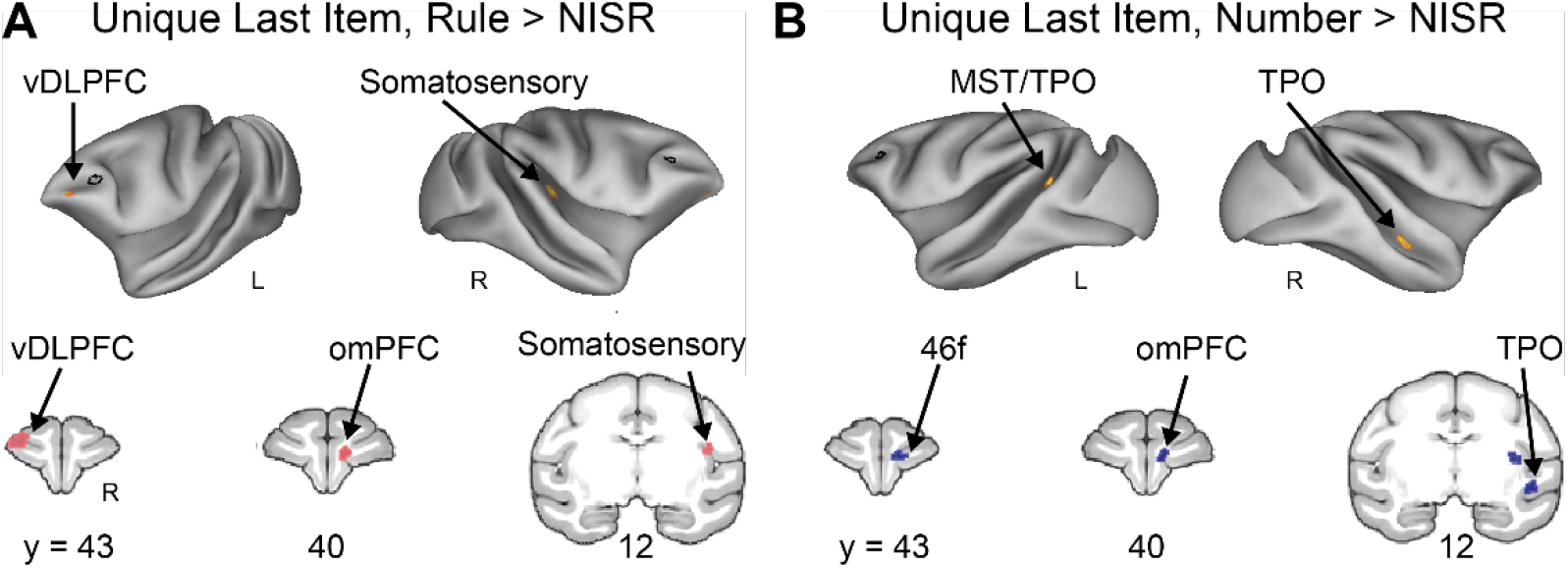
DLPFC does not show significant responses to unique last items during abstract sequence deviants. **A**. Unique Last Item, Rule Deviants > NISR (FDRc < 0.05, height p < 0.005 unc., extent = 148). **B**. Unique Last Item, Number Deviants > NISR (FDRc < 0.05, height p < 0.005 unc., extent = 82). Black outline on inflated brains indicates location of L46 or R46 (depending on the hemisphere shown) for reference. Ventral Dorsal Lateral Prefrontal Cortex (vDLPFC), Medial Superior Temporal Cortex (MST), Superior Temporal Sulcus Dorsal Bank (TPO), Orbital Medial Prefrontal Cortex, Fundus of the Dorsal Lateral Prefrontal Cortex (46f).

To determine if clusters of activation for unique ramping and unique last item were separable in the frontal cortex, we directly compared the amount of variance assigned to each cluster for both dynamics. One possibility is that, due to thresholding at the whole-brain level, there was similar activation for each dynamic across areas 46d and 46v, but that the peak, and thus the location of the thresholded cluster, differed slightly in location. To address this possibility, we created two ROIs from the clusters of significant activation in Unique Ramp, Rule Deviants > NISR contrast (area 46d, center *xyz* = −12.2, 36, 23.8 mm), and Unique Last Item, Rule Deviants > NISR contrast (area 46v, center *xyz* = −12.3, 42.9, 21.8 mm). We found a significant interaction between ROI and model in rule deviants compared to NISR (**Figure 8**; sequence type: F(1, 40) = 0.13, *p=* 0.71, η_p_^2^ = 0.003; monkey: F(2, 40) = 2.23, p = 0.12, η_p_^2^= 0.1; brain area: F(1, 40) = 0.09, p = 0.77, η_p_^2^ = 0.002; monkey x sequence type: F(2, 40) = 0.33, p = 0.72, η_p_^2^ =0.016; brain area x sequence type: F(1, 40) = 29.3, p < 0.001, ηp2 = 0.42), indicating that responses in the ramping and last item clusters were reliably different. As expected from the whole-brain contrasts, there was no significant interaction for number deviants compared to NISR (not shown; F(1, 40) = 0.796, *p* = 0.38, η_p_^2^ = 0.02). These results show that these nearby clusters in dorsal and ventral area 46 are separable in their dynamics.

**Figure 8.**
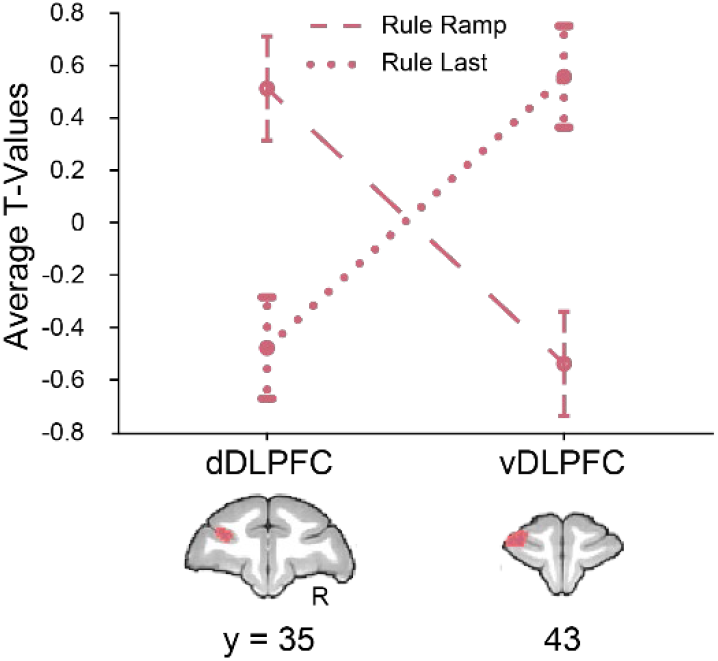
Nearby regions of DLPFC show significantly different dynamics. ROIs constructed from significant areas of activation in unique ramping (46d, dDLPFC) and unique last item (46v, vDLPFC) contrasts shown on coronal sections. T-values for the condition of interest > baseline shown. Models show a double dissociation in left area 46 during rule deviants. Error bars are 95% confidence intervals (1.96 x standard error of the within-bin mean).

## Discussion

In this study, we examined if and how monkey DLPFC (area 46) monitors abstract sequential information. We tested two main hypotheses: First, that evidence of abstract sequence monitoring would occur in the area of monkey DLPFC analogous to human RLPFC previously shown to be critical for abstract sequential tasks (Desrochers et al., 2015, 2019; McKim and Desrochers, 2022); and, second, that ramping dynamics would be associated with abstract sequence monitoring and show changes when abstract sequential structure changed. These hypotheses were tested in a no-report sequence viewing task that allowed the isolation of dynamics associated with sequence monitoring from other potential confounds such as motor preparation. We found evidence to support both hypotheses. Right area 46 responded to abstract sequence changes in both rule and number. Interestingly, left area 46 also responded to changes in abstract sequential rules, but with ramping dynamics that were similar to those observed in humans and separable from an increase only at the last item. These results suggest that a specific subregion of monkey DLPFC is specialized for monitoring general visual abstract sequential information and is a point of critical potential functional homology between human and monkey PFC during higher-level cognitive function.

The activation patterns found, with ramping dynamics primarily on the left and onset-based on the right, are consistent with prior findings. Though not explicitly tested, in humans ramping was observed preferentially on the left during abstract sequential tasks (Desrochers et al., 2015, 2019), Experiment 1) in contrast to bilateral ramping activation observed in tasks where sequences were based on stimulus identity (i.e., ordered visual items; (Desrochers et al., 2019), Experiment 2; (McKim and Desrochers, 2022). These human results are also consistent with the long-standing literature that emphasizes abstract cognitive functions in the left hemisphere of humans (e.g., (Badre and D’Esposito, 2007; Bunge et al., 2009; Wendelken et al., 2012), with the most famous such example being language (Broca, 1861; Milner, 1971; Petrides, 2013). Ramping activity has been observed in human frontal cortex during sequential language information processing (i.e., sentence comprehension) with electrocorticography (ECoG) (Fedorenko et al., 2016). While generally consistent with prior work, future studies of the distinct left and right hemisphere activation dynamics will be needed to determine their underlying drivers and their potential cognitive import. In particular, the present findings suggest that neural dynamics underlying DLPFC sequence monitoring may be distinct to each hemisphere. The monkey fMRI findings described here can provide a guide for such recordings. In sum, our results raise the possibility that the representation in and contribution of DLPFC to abstract sequence monitoring is lateralized in monkeys, and provide a road map to future studies.

An advantage of fMRI is the whole brain view that is not afforded by typical electrophysiological techniques in macaques. This view leads to potential insights about functional organization of brain areas without the limitations of a recording chamber (Milham et al., 2022). For example, in recent literature, fMRI has enabled the mapping of projections to and from the PFC with a level of specificity and across distances not previously possible on this scale (Xu et al., 2022). This work found an overlap in topographically organized high-level visual maps from the dorsal and ventral streams in primate lateral prefrontal cortex. These results raise the possibility that the localization of abstract visual sequence monitoring in DLPFC results from its position near the apex of highly organized visuo-spatial maps. The overarching organization of more cognitive processes in monkey frontal cortex has remained more elusive (Hutchison and Everling, 2014; Neubert et al., 2014, 2014; Saleem et al., 2014), and the results presented here represent a critical step forward in understanding their topography.

We observed similarities and differences to a previous study using a similar auditory task (Wang et al., 2015) that may reflect the modality employed and the capacity for generalization. Areas of the brain that responded to deviant sequences in both the current visual and prior auditory studies may be modality general. These areas included premotor cortex, caudate nucleus, and the auditory cortex rostromedial belt. In contrast, brain areas that uniquely responded to deviants in the auditory or this visual task may be modality specific for abstract sequence changes. For example, deviant responses in ventral LPFC and superior temporal sulcus were unique to the auditory study. In this visual task, deviant responses in DLPFC, visual cortex, and mPFC were observed that were not observed in the auditory task. Though we cannot draw strong conclusions without direct comparison between the modalities, these results suggest that networks of brain areas that partially overlap may constitute abstract sequence tracking across modalities. One further important difference in the studies is that in the auditory study, there was a lack of overlap between areas that respond to rule and number deviants in the monkey (in contrast to the human). Here, we observed overlap in these responses in area 46, suggesting a higher level of visual integration in the monkey. The question of sensory domain generality and integration remains open to further investigation.

The results observed here in this no-report abstract sequence viewing task in monkeys are similar to those observed in humans during sequential tasks in important ways. First, ramping activation was observed in similar regions in the frontal cortex in the monkey (area 46) and human (RLPFC). Other similar areas included visual cortex, putamen, and pre-motor cortex (Desrochers et al., 2015, 2019; McKim and Desrochers, 2022). Though the current experiment was only designed to detect the presence of ramping in relation to abstract sequences, by analogy, the function may be similar in the lateral prefrontal cortex across species. Ramping activity has been ascribed to many possible functions across species, including accumulating evidence (de Lange et al., 2010; Krueger et al., 2017; Darriba and Waszak, 2018; Lin et al., 2020), keeping time (Nobre et al., 2007; Berdyyeva and Olson, 2011; Cueva et al., 2020), reward anticipation (Roesch and Olson, 2007; Horst and Laubach, 2013; Chiew et al., 2016; Falcone et al., 2019; McKim and Desrochers, 2022) and monitoring sequence position (Desrochers et al., 2015, 2019). These possibilities are not mutually exclusive, as recent evidence in humans suggests that reward anticipation and sequence information may be present simultaneously in this signal (McKim and Desrochers, 2022). The current experiment identified ramping in the DLPFC as being sequence related, but did not examine other potential influences and therefore remains an open avenue of future inquiry. Similarly, other brain areas that display ramping activation remain open for investigation.

Though this study bears resemblance to a field of literature using statistical learning paradigms, the majority of those studies use tasks that rely on the identity of the stimuli themselves. Importantly, this study is distinct from most statistical learning studies because the identity of the stimulus alone cannot predict the following item (i.e., knowing the current fractal is the green one does not determine the next stimulus without also having sequential rule information). A subset of work in the infant learning literature examines statistical learning that is not dependent on stimulus identity (i.e., “artificial grammar”), but mostly auditory tasks were used (e.g., (Saffran et al., 1996)). To our knowledge, only one behavioral study examined violations to a visual (and auditory) artificial grammar where sequences of different lengths are constructed according to set transition probabilities (Milne et al., 2018). While the findings were important because violations were detected similarly in monkeys and humans, there were no neural data presented. Further, while transition probabilities created the sequences, there was not a set “rule” by which they were constructed. Therefore, the present study is unique in examining neural responses to abstract visual sequences.

This study contained the following limitations. First, the timing of the stimuli in the current design did not allow examination of dynamics of individual sequence items, only across the sequence as a whole. In a single experiment, it was not feasible to separate each sequential item by the time required to model each separately in an event-related design. Therefore, future work will aim to examine the dynamics of individual sequential items in greater detail. Second, while the no-report task allowed the elimination of motor preparatory confounds, it did not allow for direct correlation with behavioral performance. Although the observed signals will potentially also underlie tasks that require responses, this assertion remains to be tested and the present study is an important foundation for further experiments. Third, we have focused here on the DLPFC because although its importance in cognitive processes in monkeys has been established, its response to visual abstract sequences and potential correspondence to dynamics in humans remained unknown. The DLPFC is part of a network of areas active in this task, and although they are outside the scope of the current experiment, they remain an important avenue of future research.

In summary, we provide evidence that a specific subregion of monkey DLPFC monitors abstract visual sequences and generalizes across different sequence violations (number and rule). Further, sequence related ramping dynamics were also observed in DLPFC. Importantly, this region is possibly analogous to human RLPFC, where necessary sequence-related ramping signals have been identified in the past. These results suggest functional homology across the species as to where and how more general abstract visual sequential information is represented in the brain. These findings, in turn, inform future models of how abstract sequential information is represented during more complex behaviors across species.

## Acknowledgements

We thank Matthew Maestri for his assistance with animal training and data collection. We would also like to thank Dr. Michael Worden, Lynn Fanella, Fabienne McEleney and Brown University’s MRI Facilities staff for their support and guidance throughout this project. We also thank Dr. David Sheinberg, Dr. Amitai Shenhav, Dr. Katherine Conen, and Dr. Diana Burk for their continued support throughout this project and preparation of this publication, as well as members of both the Sheinberg and the Desrochers Labs for many helpful discussions and contributions. This study was supported by the NSF-EPSCoR Neural Basis of Attention 1632738 (N.Y.R, T.M.D.), the NIGMS-NIH Initiative to Maximize Student Development IMSD R25GM083270 (N.Y.R.), the NIH-NIGMS COBRE P20GM103645-07, and the Carney Institute for Brain Science Innovation Award (T.M.D.). Part of this research was conducted using computational resources and services at the Center for Computation and Visualization, Brown University, S10 OD016366.

## Extended Data

**Figure 3-1.**
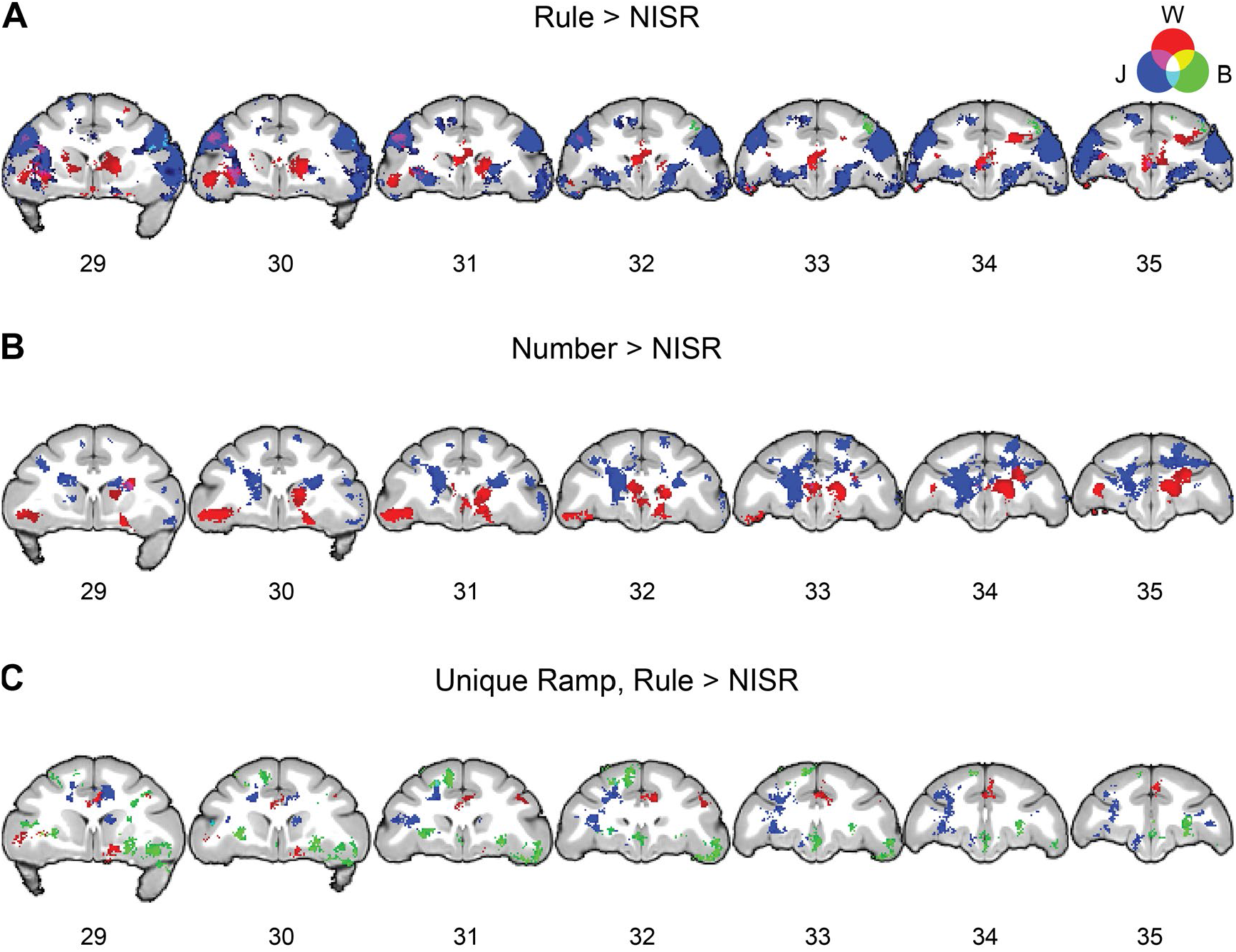
Overlaid individual monkey contrasts for monkeys W, J, and B. Three second level contrasts, one for each monkey, were created using only data bins that contained runs from each animal (W = 9, J = 6, B = 7 bins each). As described in Materials and Methods, each bin contained approximately 10 runs. A liberal height threshold of p < 0.05 was chosen for illustrative purposes before applying and extent (extents listed for each monkey by contrast) and false discovery rate (FDR) error cluster corrected for multiple comparisons to p < 0.05. Slice number in the y direction is listed under each coronal section. **A**. Voxel wise contrast of Rule Deviants > New Items, Same Rule (NISR) (W extent = 560, J = 1296, B = 623 voxels). **B**. Voxel wise contrast of Number Deviants > NISR (W extent = 772, J = 750, B = 667 voxels). **C**. Voxel wise contrast of Unique Ramp, Rule Deviants > NISR (W extent = 560, J = 630, B = 581 voxels).

**Figure 4-1.**
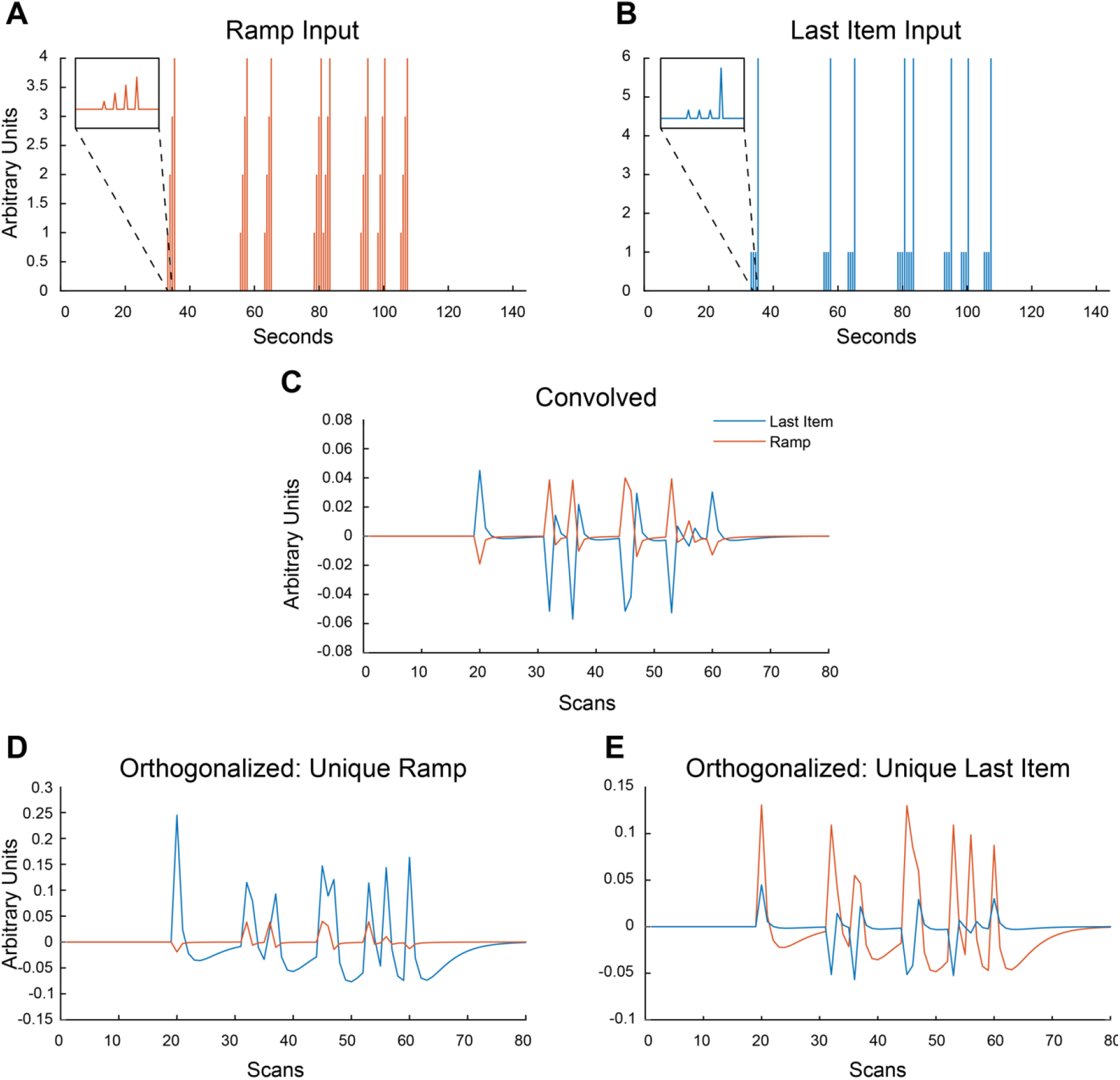
Example Ramp and Last Item regressors through the orthogonalization process. Note that SPM first creates regressors from onsets (in seconds, shown in **A**.) in samples using higher resolution to be convolved, and then they are down-sampled before being orthogonalized and entered in the GLM.

